# Populations of the Australian saltmarsh mosquito *Aedes vigilax* vary between panmixia and temporally stable local genetic structure

**DOI:** 10.1101/2025.01.05.631387

**Authors:** Thomas L Schmidt, Nancy Endersby-Harshman, Toby Mills, Rahul Rane, Gunjan Pandey, Chris Hardy, Leon Court, Cameron Webb, Brendan Trewin, Brett Neilan, Ary A Hoffmann

## Abstract

Pest management programs can operate more effectively when movement patterns of target species are known. As individual insects are difficult to track, genomic data can instead be used to infer movement patterns based on pest population structure and connectivity. These data can also provide critical information about cryptic taxa relevant to management. Here we present the first genomic investigation of *Aedes vigilax*, the Australian saltmarsh mosquito, a major arbovirus vector across Australasia. We used a ddRAD pool-seq approach and a draft genome assembly to investigate genetic variation in 60 *Ae. vigilax* pools from across Australia but with a focus on urban Newcastle and Sydney, NSW. There was strong genetic structure between samples from the west and east coasts of Australia, and additional structure that differentiated east coast populations. Within Newcastle and Sydney, contrasting patterns of genetic structure were evident. In Newcastle, there was no differentiation among subregions up to 60 km apart. In Sydney, samples from one urban subregion were differentiated from others < 3 km apart, and this structure was stable across sampling years. Heterozygosity and Tajima’s D indicated no bottlenecks in Newcastle or Sydney populations, suggesting this structure represents a gene flow barrier. Nuclear differentiation patterns contrast with previous mtDNA data indicating two COI clades in the east coast, one of which was also present in Western Australia. The panmixia over 60 km across the Newcastle region corroborates previous field observations of high dispersal capacity in this mosquito. These findings indicate specific challenges that may hinder local suppression strategies for this species.

## Introduction

Traditional approaches to the management of pests like mosquitoes involving widespread pesticide applications face many well-known challenges such as environmental damage and the evolution of pesticide resistance (Martelli et al. 2022; Schmidt et al. 2024a). On the other hand, the development and operationalisation of novel approaches, including targeted local pesticide applications, the release of *Wolbachia* and sterile males for population suppression, and the release of modified pests expressing advantageous traits, can be difficult when there is a lack of entomological knowledge of target populations (Benelli 2015; Fernandes et al. 2018). Genomic research can provide some of this fundamental knowledge and supplement ecological studies, which has led to a rapid proliferation of population genomic studies of pests aiming to inform species- or population-specific integrated pest management strategies (Xu et al. 2022; Urvois et al. 2022; McCulloch et al. 2023; Schmidt et al. 2023b; Raszick et al. 2024). For example, genomic studies can detect cryptic species that may go unobserved through morphological identification, and can provide information on hybridisation and introgression not always evident from lower-powered molecular markers such as microsatellites or mtDNA (Valencia-Montoya et al. 2020; Small et al. 2023; Caputo et al. 2024). Once species relationships are established, genomics can be used to infer patterns of connectivity and barriers among pest populations at increasingly fine scales, including within targeted urban control regions and surrounding areas (Schmidt et al. 2023a; Paris et al. 2023; Booth 2024). These spatial patterns can be contrasted with temporal patterns of population structure, where genomic data can help distinguish overwintering populations from those receiving new recruits each season, and potentially determine the source of these recruits (Hereward et al. 2020; Chen et al. 2021; Raszick et al. 2024). Finally, genomic patterns within and across populations can reveal potential past bottlenecks caused by the invasion process or subsequent population suppression (Dogantzis et al. 2024; Hagan et al. 2024), which can indicate how the population may evolve in response to new selection pressures such as from insecticides or other control operations (Charlesworth 2009).

Until the present decade, high sequencing costs mostly restricted pest genomic studies to major international vectors of human health concern (e.g. *Anopheles* and *Aedes* mosquitoes (Small et al. 2023; Schmidt et al. 2024a)) and international herbivorous pests (e.g. *Helicoverpa* and *Plutella* moths (Chen et al. 2021; Jin et al. 2023)). Current sequencing costs are far lower, allowing for other species to be analysed, including those that are widespread (Comeault et al. 2020; Xu et al. 2022) but also those of more local importance (Raszick et al. 2020; Thia et al. 2023). This latter group includes many mosquito species without the global distributions of species like *Aedes aegypti*, *Anopheles stephensi*, and *Culex quinquefasciatus*, but which may nevertheless be important local vectors of pathogens of human health concern (Henderson et al. 2023; Paris et al. 2023).

For Culicine mosquitoes (e.g. *Aedes* and *Culex* spp.), genomics research has been challenged by the large and repetitive genomes of these species as well as high residual heterozygosity prohibiting accurate haploid assemblies (Deng et al. 2024). Currently, only ∼5 of >3000 Culicine species have reference genome assemblies available. Draft genome assemblies can nevertheless be sufficient for answering many key questions in pest control (Paris et al. 2023), and full genomes of Culicine mosquitoes are now emerging more frequently (e.g. Ryazansky et al. 2024), owing in part to newer approaches using single mosquitoes for complete genome assemblies or long read methodologies.

While Australia is generally free of internationally significant mosquito-borne diseases (e.g. dengue, malaria), there is a suite of arboviruses of human health concern. Ross River virus (RRV) infects approximately 5,000 people every year, and although the disease is not fatal the symptoms can be severely debilitating (Claflin and Webb 2015) The incidence of RRV activity in coastal regions of Australia has increased in rural and metropolitan areas (Jansen et al. 2019) and drawn the attention of health authorities (Skinner et al. 2020; Murphy et al. 2020). The public health concerns associated with RRV have increased interest in management of the mosquito vectors of these viruses.

This study presents a draft genome assembly and a population genomic analysis of the Culicine mosquito pest, *Ae. vigilax*, a mosquito of significant pest and public health concern in coastal regions of Australia (Webb et al. 2016). This mosquito is found throughout Australia and in some other Indo-Pacific countries (Lee 1980; Puslednik et al. 2012) and its abundance is expected to increase under recent climate change and sea level rise (Staples et al. 2024). *Aedes vigilax* is common in estuarine ecosystems and closely associated with tidally influenced saltmarsh and mangrove habitats, and also occurs sporadically away from the coast. It is well established as a severe nuisance-biting pest and major vector of RRV and Barmah Forest virus (BFV). As a result, mosquito control programs are in place to target this species by many local authorities (Russell and Kay 2008). However, other regions of coastal Australia do not have comprehensive mosquito control programs in place, and with growing residential and recreational developments in many areas a focus has been on how best to manage *Ae. vigilax* and the public health threats it poses. The Newcastle region of New South Wales, Australia (Fig. 1), has been one such area where traditional mosquito control faces challenges due to cost, multi-agency land management, and the presence of ecologically sensitive coastal wetlands. These challenges have prompted interest in developing strategies to suppress or replace this population using a rear-and-release program involving endosymbiotic *Wolbachia* bacteria, similar to methods used in *Ae. aegypti* (Hoffmann et al. 2011; Beebe et al. 2021) and *Ae. albopictus* (Zheng et al. 2019).

**Fig. 1.**
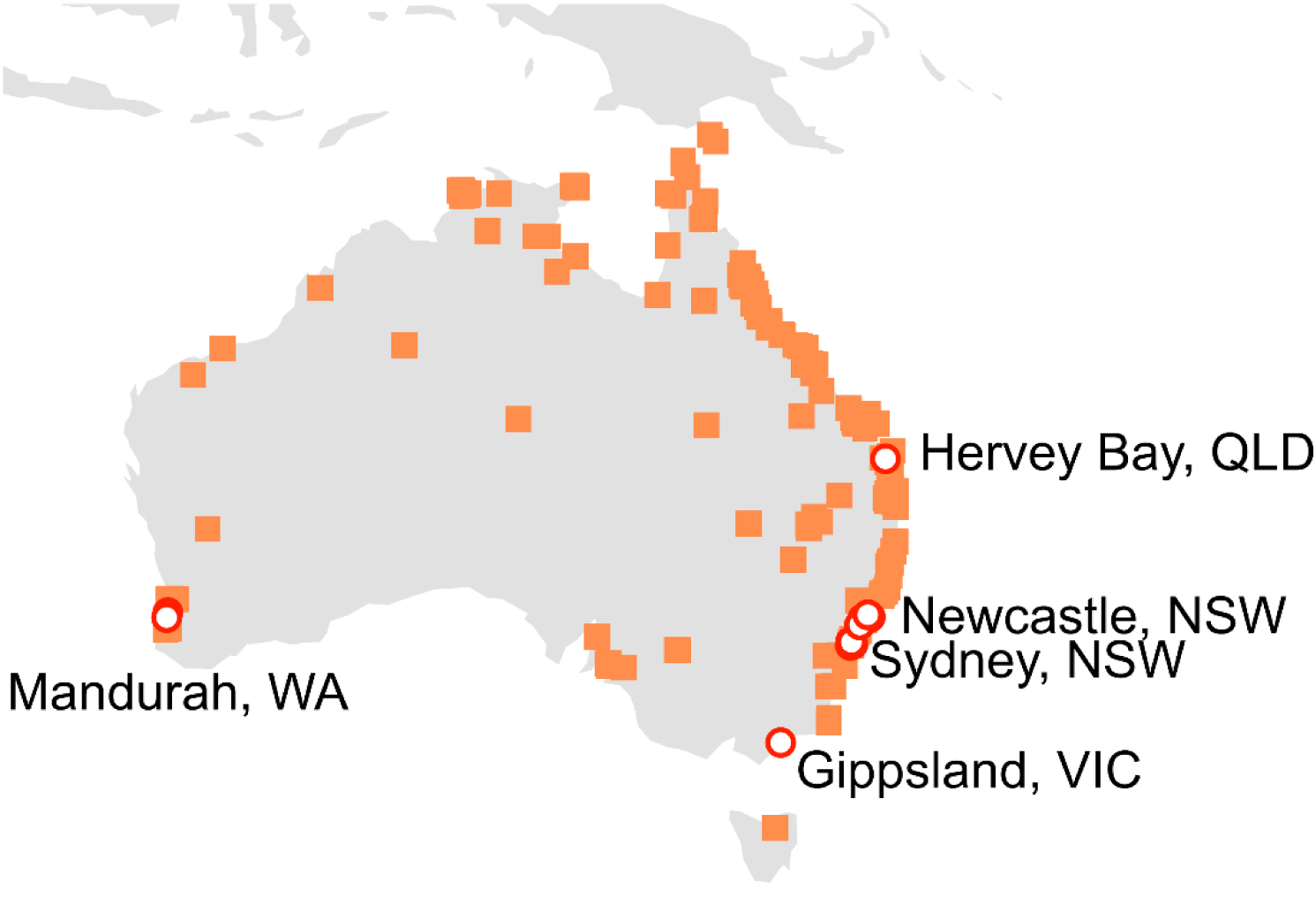
Locations in Australia where *Aedes vigilax* has been recorded since 1950 (Atlas of Living Australia; squares), and locations where samples were taken for this study (circles).

Despite the importance of *Ae. vigilax*, many gaps remain in our understanding of this species’ biology across its known distribution and around Newcastle region specifically, including knowledge critical to the effectiveness of novel management programs. First, it is unclear whether *Ae. vigilax* is a species complex, as found in other local mosquitoes including the sympatrically distributed Australian mosquito, *Ae. notoscriptus* (Endersby et al. 2013). A molecular genetic study based mainly on mtDNA variation found two sympatrically distributed clades of *Ae. vigilax* on Australia’s east coast with 4.2% divergence at the COI gene (Puslednik et al. 2012), which may represent two cryptic subtaxa of *Ae. vigilax;* only one of these occurred in Western Australia. However, divergence in two nuclear sequences was less clearly associated with these putative subtaxa (Puslednik et al. 2012). Second, it is unknown whether populations of *Ae. vigilax* in Newcastle and other regions represent single panmictic populations or structured populations with locally restricted gene flow. Fine-scale genetic structure reflecting restricted urban gene flow occurs in other *Aedes* including *Ae. aegypti* (Schmidt et al. 2023a) and *Ae. albopictus* (Schmidt et al. 2017b; Yeo et al. 2023) but weaker structure has been recorded in a stronger-dispersing species, *Ae. notoscriptus* (Paris et al. 2023). *Aedes vigilax* is thought to be one of the most widely dispersing mosquitoes in Australia, as it has been recorded moving >3 km to and from offshore islands (Johnson et al. 2020), between widespread wetlands (Chapman et al. 1999), and across urban wetlands (Webb and Russell 2019). Anecdotal evidence suggests the mosquito may disperse more than 20km from immature habitats (Lee 1980). Third, it is unclear whether spatial genetic structure is temporally stable, such as might be expected if recruitment over seasons is locally sourced, or whether recruitment is from distant sources. Finally, while the species is widely distributed in Australasia (Fig. 1), it is not yet clear which populations constitute the original invasive range; COI data suggest initial separation between eastern and western populations with secondary contact, but data from two nuclear sequences are less clear on this scenario (Puslednik et al. 2012).

These four issues – species complexes, spatial structure, temporal structure, and population history – are all critical to understanding the biology of *Ae. vigilax* and developing targeted control programs, particularly for rear-and-release strategies. Unrecognised species complexes create general difficulties for control but can be catastrophic for rear-and-release strategies as these require interbreeding between lab-reared mosquitoes and field mosquitoes to achieve population suppression or replacement. Mosquito releases of a different subspecies to those of the target population will have little to no impact on these outcomes. Spatial patterns of genetic structure indicate regions of restricted gene flow, which allows for a single large population to be reclassified into a series of subpopulations that can be specifically targeted for mosquito releases and other control methods (Schmidt 2024). Successful rear-and-release interventions have been undertaken in various isolated towns in Queensland, Australia (Hoffmann et al. 2011; Beebe et al. 2021), with more mixed results in some larger urban areas (Nazni et al. 2019; Santos et al. 2022). Patterns of temporal structure and seasonal recruitment can indicate how an intervention may play out over successive years, where populations having only local recruitment will remain controlled in subsequent years, while those receiving external recruits may require repeated interventions each season. Finally, large mosquito populations will require proportionately larger numbers of released mosquitoes to achieve control targets, though large populations can produce more stable outcomes for strategies aiming to permanently drive *Wolbachia* bacteria into a wild population (Schmidt et al. 2017a).

Here we use a spatially continuous and temporally cross-sectional sampling design to investigate species complexes, spatial structure, temporal structure, and population history in *Ae. vigilax.* We focus on two urban populations from the Newcastle and Sydney regions of New South Wales, Australia, with additional samples from three other Australian states and with a draft genome assembly to guide the analysis. We found no evidence of cryptic species, but observed patterns of genetic structure within the two urban populations that were surprisingly heterogeneous. Our findings will directly inform future management strategies for *Ae. vigilax* populations across Australia and provide a range of general insights for urban mosquito management worldwide.

## Methods

### Draft genome sequencing and assembly

Samples of *Ae. vigilax* were collected from Brisbane, Queensland, Australia, and cultured in the laboratory. High molecular weight DNA was extracted using the Qiagen Genomic Tip 20/G kit (Qiagen, Cat#: 10223) and the Qiagen Buffer Set (Qiagen, Cat#: 19060). The ‘user-developed protocol’ for ‘mosquitoes and other insects’ supplied by Qiagen was followed, except EB buffer was used to dissolve the purified genomic DNA instead of Tris-EDTA buffer. For Illumina short read sequencing, Illumina PCR-based libraries were constructed as per the manufacturer’s protocol and sequenced to approximately 60× coverage on an Illumina NovaSeq 6000 sequencer, S4 flow cell lane (2x150bp PE). For long read sequencing, a standard PacBio library was prepared following the manufacturer’s instructions. The preparation involved shearing the DNA, end-repair, and ligation of SMRTbell adapters. This library was sequenced to 50× coverage using the Pacbio Sequel IIe sequencer with an average read length of 15 kb.

Illumina data were cleaned and adapter sequences removed from the resulting reads using Trim-Galore (v 0.6.6). PacBio data were error corrected to increase average Q-scores using Ratatosk (Holley et al. 2021). The genome was assembled from the longest error-corrected 30× reads using the Raven assembler (Vaser and Šikić 2021), and further refined through the following: three rounds of polishing with long reads using Racon (Vaser et al. 2017); three additional rounds of polishing with short reads using Racon; and six rounds of Polca polishing using the Masurca package (Zimin et al. 2013).

Final genome assembly statistics are listed in Table 1. The assembly is accessible through the CSIRO Data Access Portal (Rane et al. 2024).

**Table 1.** Genome assembly statistics.

|  |  |
| --- | --- |
| Total Number of Contigs | 29,254 |
| Total Genome Size (bp) | 1,264,995,562 |
| Contig N50 (bp) | 50,356 |
| Contig N50 (n) | 8,264 |

### Field samples, library preparation, and sequencing

Adult *Aedes vigilax* were collected from locations across the Newcastle and Sydney regions in New South Wales (NSW), Australia. Samples were collected in 2021 and 2022, in March and April, which represents the end of the mosquito season in each year. Adult mosquito sampling locations were determined by assessing actual and potential immature habitats (i.e. saltmarsh, mangrove, sedgeland vegetation communities) and associated adult refuge locations (i.e. bushland). The Newcastle region samples were collected in 2021, while the Sydney region samples were collected across 2021 and 2022, including two regions that were sampled in both years: Duck River (Silverwater, western Sydney), and Sydney Olympic Park. Additional samples were collected from other states: Western Australia (Mandurah region, WA), Victoria (Gippsland, VIC), and Queensland (Hervey Bay, QLD) (Fig. 1). Adult mosquitoes were sampled using carbon dioxide baited Encephalitis Virus Surveillance (EVS) traps (Rohe and Fall 1979). Mosquitoes were identified to species using pictorial guides (Webb et al. 2016) and taxonomic keys (Russell 1993). Only specimens that were confidently confirmed through morphological identification as *Ae. vigilax* were retained for further analysis.

We used these samples to construct double digest RAD sequencing (ddRADseq) libraries of *Ae. vigilax* DNA pooled across 50 individuals. Each pool represented a sampling location and time. Only the head was used from each individual, to normalise the quantity of DNA across the 50 individuals. Two pools had fewer than 50 individuals due to sampling constraints; these were Rockdale, NSW (36 individuals) and one of the two pools in Dora Creek, Newcastle region, NSW (49 individuals). Full sample details of pools are in Table S1. Samples listed as “(aggregate)” are those in which trap collections were aggregated across 2-3 time points to make up the full 50 individuals; these time points were always within a two-week range.

Genomic DNA was extracted from each pool of mosquito heads using the High Pure PCR Template Preparation Kit (Roche Diagnostics GmbH, Mannheim Germany). Intense pigmentation from 100 mosquito compound eyes in each sample necessitated extra centrifuge steps and pipetting against a light source to identify and remove exoskeleton debris. An RNase step was not included in the protocol and final elution of DNA was made in 200 µL of Elution Buffer (10 mM Tris-HCl, pH 8.5 (+25°C)).

Genomic DNA libraries were constructed using the ddRADseq method (Peterson et al. 2012) as modified (Rašić et al. 2014) with further refinement to allow sequencing of multiple small libraries in a single sequencing lane (Schmidt et al. 2023a). Three genomic libraries were constructed, each containing DNA from 21 pools of mosquito heads. For each pool, 300 ng of genomic DNA were digested with restriction enzymes, MluCI and NlaIII (9 units each) in NEB CutSmart buffer (4.5 µL) (New England Biolabs, Ipswich, MA, USA). The volume of each pool’s digestion reaction was made up to 45 µL with PCR-grade water and digestion took place for 3 h at 37°C. After digestion, the DNA was cleaned with 1.5x Sera-Mag Magnetic Carboxylate-Modified Particles (Life Sciences IP Holdings Corporation, Washington, USA). To allow identification of individual pools, Illumina P1 and P2 adapters with barcode modifications (2.0 µL of 1.8 µM each) were joined to the cleaned DNA fragments in an overnight ligation at 16°C using T4 DNA ligase (1100 units) and T4 Buffer (4.5 µL) (New England Biolabs, Ipswich, MA, USA). DNA was normalised to 35 ng with PCR-grade water to a volume of 35.95 µL before heat denaturation at 65°C. Fragments were cleaned with 1.5x Sera-Mag Magnetic Carboxylate-Modified Particles after the ligation reaction.

Size selection to 300 – 450 bp of the resulting fragments was undertaken using a Pippin Prep instrument and a 2% agarose with ethidium bromide, 100 – 600 bp, Marker B gel cassette (Sage Science Inc., Beverly, MA, USA). After size selection, 1 µL of DNA was amplified by PCR using 5 µL of Phusion High Fidelity 2x Master mix (Thermo Scientific Inc., Waltham, MA USA) and 2 µL of 10 µM standard Illumina P1 and P2 primers. PCR cycling conditions were 98°C denaturation for 30 s followed by 12 cycles of 98°C (denaturation) for 10 s, 65°C (annealment) for 30 s and 72°C (extension) for 70 s with a final extension at 72°C for 5 min. Each library was amplified with the same Illumina forward primer (PCR1_D0501_TATAGCCT) and a different Illumina barcoded reverse primer to allow sequencing in a single lane (Library A: 2-1: PCR2_D705_ATTCAGAA, Library B: 2-2: PCR2_D706_GAATTCGT, Library C: 2-3: PCR2_D707_CTGAAGCT). PCR amplicons were cleaned using 1.5x Sera-Mag Magnetic Carboxylate-Modified Particles before being quantified and sent for sequencing to Novogene Co., Ltd. (Beijing, China).

Libraries were sequenced on a NovaSeq 6000, to obtain 150 bp paired-end reads, with an average of 67.2 Gbp data per library. Three pools were omitted due to insufficient reads. The final filtered dataset consisted of 38 pools from Newcastle region, 18 pools from Sydney region, 2 pools from WA, 1 pool from VIC, and 1 pool from QLD.

### Sequence alignment, genotyping, and downsampling

We aligned sequences to the *Ae. vigilax* genome assembly using bowtie2 with “--very-sensitive” alignment settings. The proportion of reads that successfully align to a reference assembly tends to be higher for genetically similar samples and decreases with genetic distance (Thorburn et al. 2023). For highly differentiated samples, such as of different cryptic species, read rates may be far lower. Accordingly, we expected alignment rates would be much lower for any cryptic subspecies of *Ae. vigilax* included in the study.

Our initial analyses identified a positive relationship between the number of aligned reads for a pool and the autosomal heterozygosity of that pool (see below section). Accordingly, we used samtools view v1.16 (Danecek et al. 2021) to downsample reads in each pool so that all pools had equal read numbers, using the “-s” command. There was no relationship between heterozygosity and read number after downsampling (R^2^ = 0.001, F_1,58_ = 0.050, P = 0.82). These downsampled files were used for all subsequent analyses.

### Spatial and temporal genetic structure

We investigated the genetic relationships between pools using TreeMix (Pickrell and Pritchard 2012), pairwise F_ST_, and principal components analysis (PCA). These analyses were used to detect highly differentiated populations that could represent cryptic taxa, as well as for characterising how genetic differentiation among pools varied across space and across the two sampling years.

First, we used FreeBayes v1.3.2 (Garrison and Marth 2012) to genotype all 60 bam files simultaneously, using standard filters (minimum mapping quality = 30, minimum base quality = 20). We then used vcftools (Danecek et al. 2011) to filter the resulting VCF file to filter sites with any missing genotypes, with read depth < 15× in any individual, and with minor allele frequency across all pools of < 0.05. This left 9331 filtered SNPs.

We ran TreeMix to build maximum likelihood trees for the 60 pools, using the -k option to analyse blocks of 50 SNPs to limit issues from analysing linked sites. We tested the effects of adding migration edges to the tree by running TreeMix for each of 0, 1, 2, … 7 migration edges, and assessing the second-order rate of change in likelihood from adding each additional edge across ten replicates (Fitak 2021).

We estimated pairwise F_ST_ using the R package “poolfstat” (function “compute.pairwiseFST”). Scores were transformed into linear F_ST_ (F_ST_ /(1 – F_ST_) and Mantel tests were used to assess isolation by distance between linear F_ST_ and the natural logarithm of pairwise geographic distance. Mantel tests were run in R package “ecodist” (function “mantel”) using 10,000 permutations. For PCA, we used “poolfstat” to estimate allele frequencies from the read depths of each allele, then used the R function “prcomp” to run a PCA on the matrix of allele frequencies. We visualised PCA results by interpolating them with ordinary Kriging, setting an exponential semivariogram model and a search radius that included all points. This was run in ArcGIS Pro v3.1 (https://www.esri.com/en-us/arcgis/products/arcgis-pro/overview).

### Variation within individual pools

We investigated population demography using pool-specific estimates of autosomal heterozygosity and Tajima’s D (Tajima 1989), and inferred long-term effective population size in Sydney and Newcastle using the nucleotide diversity (π) across all pools in each region. Heterozygosity and Tajima’s D are standard metrics of genetic variation which are expected to change in populations that have experienced bottlenecks, with heterozygosity decreasing and Tajima’s D becoming more positive. Effective population size is a parameter representing a population’s capacity to evolve by natural selection as well as the required number of mosquitoes for rear-and-release programs, with larger values indicating a greater capacity to evolve and a larger number of mosquitoes being required.

To estimate autosomal heterozygosity, we reprocessed and refiltered each pool of individuals in alignment with protocols developed for estimating autosomal heterozygosity in individuals (Schmidt et al. 2024b). Notably, these protocols help evade issues arising when data are already filtered for polymorphism (i.e. SNP data) (Schmidt et al. 2021). By processing pools individually, these protocols also allow data to be filtered with high minimum read depth cutoffs (50×) and with all sites with missing genotypes omitted, while still retaining large quantities of sequence data for parameter estimation (Schmidt et al. 2024b). For each pool, we applied FreeBayes v1.3.2 (Garrison and Marth 2012) to call genotypes from the aligned bam file, using standard filters (minimum mapping quality = 30, minimum base quality = 20, minimum supporting allele qsum = 0, genotype variant threshold = 0), excluding unobserved genotypes but retaining monomorphic loci. The output VCF file was sorted and filtered with bcftools v1.16 (Danecek et al. 2021) to remove all missing genotypes, genotypes with less than 50× read depth, and genotypes with spanning deletions. We estimated heterozygosity at each site using the read depths of the reference and alternate alleles, where the ratio of the read depth relative to total depth provided an estimate of allele frequency.

To estimate Tajima’s D, we used samtools mpileup v1.16 (Danecek et al. 2021) to build pileup outputs for each individual pool, then applied Popoolation v1.2.2 (Kofler et al. 2011) to estimate Tajima’s D in non-overlapping windows of 10 kbp, setting a minimum base quality of 30, and minimum fraction covered of 0.001 (as ddRADseq data are sparse), a minimum coverage of 10 and a maximum coverage of 300. We visualised heterozygosity and Tajima’s D results by interpolating them with ordinary Kriging, setting an exponential semi-variogram model and a search radius that included all points. This was run in ArcGIS Pro v3.1 (https://www.esri.com/en-us/arcgis/products/arcgis-pro/overview).

To estimate long-term effective population size, we used the equation: 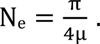 Here, N_e_ is the effective population size, π is the nucleotide diversity, and µ is the *de novo* mutation rate per generation per site (Charlesworth 2009). To estimate π across the entire sampling regions of Newcastle and Sydney, we computed a population-level metric, in which we pooled all sequence reads from all the pools in each region, and estimated π as described above for the entire set of sequence reads from the region. For µ, we applied mutation rate estimates from two studies of *Drosophila melanogaster*, namely 3e^-9^ – 4e^-9^ (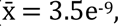 Keightley et al. 2009) and 1e^-9^ – 6.1e^-9^ (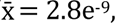 Keightley et al. 2014). A recent mutation rate estimate for *Ae. aegypti* falls within these broad ranges (4.85e^-9^; Rose et al. 2023).

## Results

### Sequence alignment and cryptic taxa

Alignment rates to the reference genome varied between 0.674 and 0.818. Comparisons involving Sydney (NSW), Newcastle (NSW), and Mandurah (WA) indicated no difference in alignment rates (Single Factor ANOVA F_2,56_ = 0.79, P = 0.46). Six pools had alignment rates lower than 0.7 (0.674 – 0.698), which were observed in the Hervey Bay (QLD) pool and five Newcastle pools. The Gippsland (VIC) pool had an alignment rate of 0.724. Based on similarity in the alignment rates, there was no evidence that any of the pools contained geographically-restricted cryptic subspecies of *Ae. vigilax* with strongly differentiated DNA sequences.

### Spatial and temporal genetic structure

Fig. 2 shows TreeMix output using zero migration edges. While the second-order rate of change in likelihood indicated that either zero or one edge was optimal, the single edge was placed inconsistently across subsequent runs of TreeMix, suggesting it had little biological relevance. The plotted tree had the same overall structure to all other trees tested by TreeMix: Western Australian (WA) pools were highly differentiated from all others, Sydney pools mostly clustered apart from Newcastle, Duck River pools clustered apart from other Sydney pools, but no structure was observed among Newcastle pools.

**Fig. 2.**
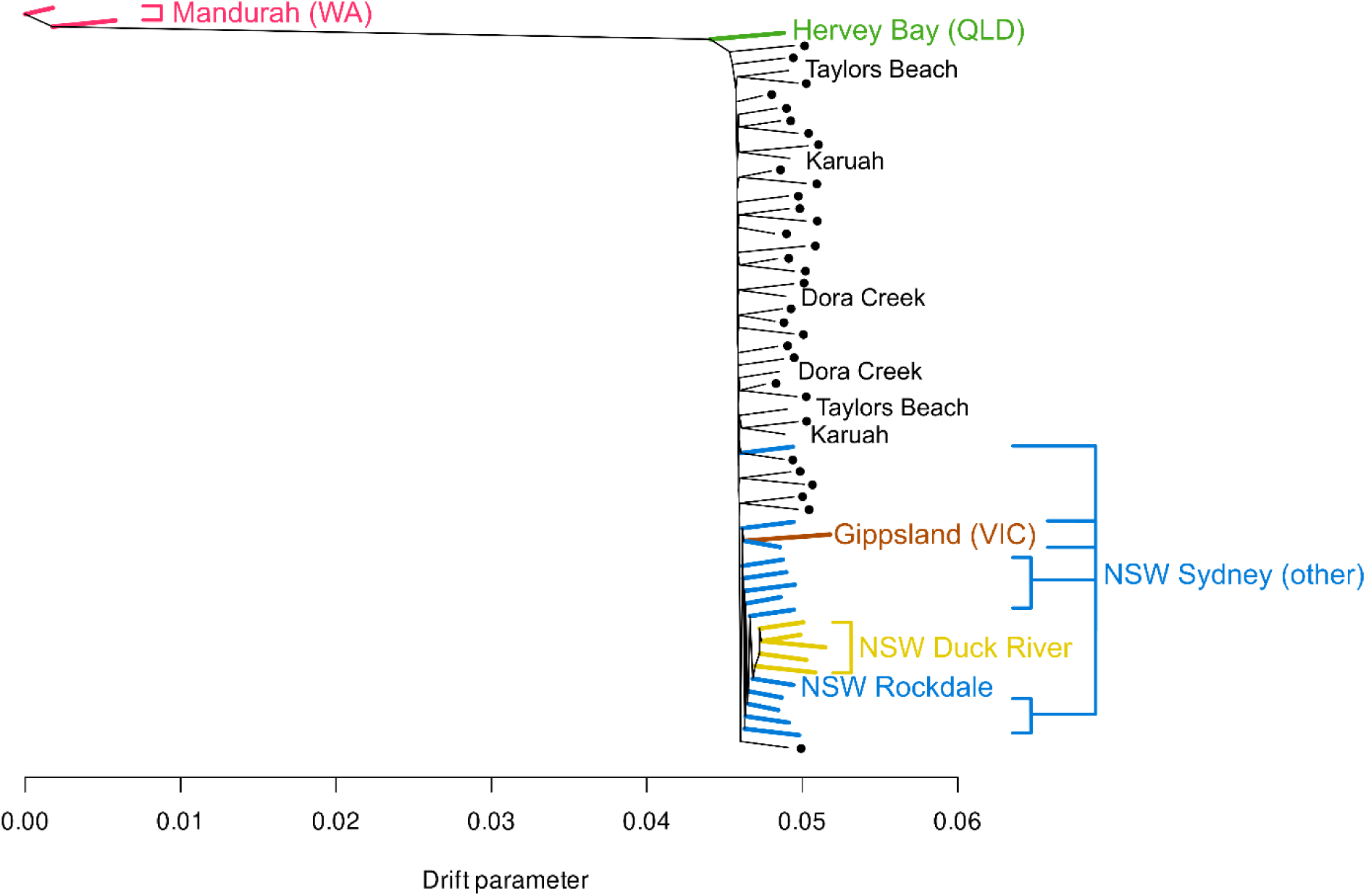
TreeMix maximum likelihood tree for all 60 pools. Colours indicate regions outside Newcastle, NSW. Newcastle pools are indicated with black dots, and peripheral Newcastle locations indicated with black text.

The lack of structure among Newcastle pools is particularly stark given there was no clustering among pairs of pools from even the most peripheral Newcastle populations of Dora Creek (south-west), Taylors Beach (northeast), and Karuah (northeast) (Fig. 3). This suggests a lack of genetic structure across the ∼60 km sampling range of Newcastle. Queensland (QLD) and Victoria (VIC) both had longer internal branches, suggesting these populations had experienced greater genetic drift than other sites, possibly due to smaller population sizes or bottlenecks during population establishment. The Hervey Bay (QLD) pool was placed between WA and NSW while the Gippsland (VIC) pool clustered with the Sydney pools (Fig. 2). Within Sydney, the clustering of Duck River samples is notable as these samples were taken over two years (2021–22), indicating these specific genetic patterns are temporally stable (Fig. 2). The most genetically similar Sydney pool to Duck River was from Rockdale, which was the most geographically isolated of all Sydney locations (Fig. 3).

**Fig. 3.**
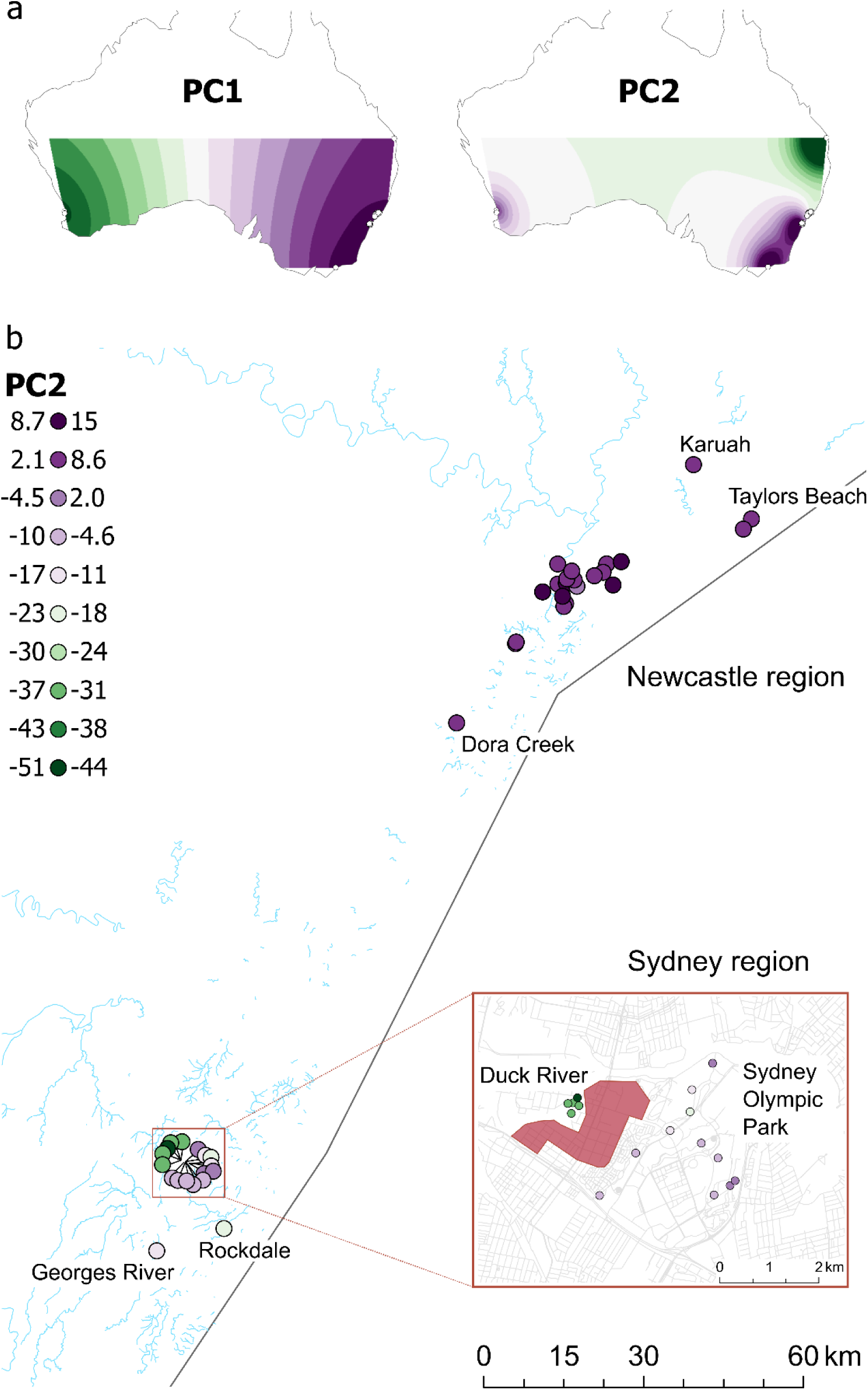
PCA results. (a) Interpolated scores for PC1 (4.98% of variation) and PC2 (2.32%). (b) Spatial variation in PC2 within NSW. Coloured circles indicate individual pools. Red polygon in inset indicates a contiguous region of industrial land use. Road and river shapefiles are from https://mapcruzin.com/.

PCA results (Fig. 3) confirmed the patterns observed in TreeMix. PC1 (4.98% of variation) differentiated Western Australia from other pools, PC2 (2.32%) differentiated Queensland from NSW, Sydney from Newcastle, and Duck River from other Sydney pools. Viewing variation on PC2 within NSW provided further insight into these patterns (Fig. 3b). Here, Newcastle samples are quite clearly differentiated from Sydney on the PC2 axis, and also in the degree of differentiation, with little structure throughout Newcastle but considerable structure within Sydney. Duck River samples were the most clearly differentiated. Additional PCs served to differentiate Gippsland (VIC) samples (PC3) and indicated further differentiation of Hervey Bay (QLD) and Duck River samples (PC4) (Fig. S1).

F_ST_ between WA pools and all other pools was very high 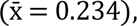 while it was much lower for all other pairs 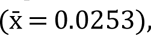 providing no evidence for any clade-based associations between east and west coast samples as suggested by the mtDNA data from Puslednik et al. (2012). After omitting WA, Mantel tests found a significant pattern of isolation by distance across the east coast samples (Fig. 4a) and between the Sydney and Newcastle samples (Fig. 4b). There was no isolation by distance within Newcastle (Fig. 4c) or within Sydney (Fig. 4d) and removing the Duck River pools made no difference to these results.

**Fig. 4.**
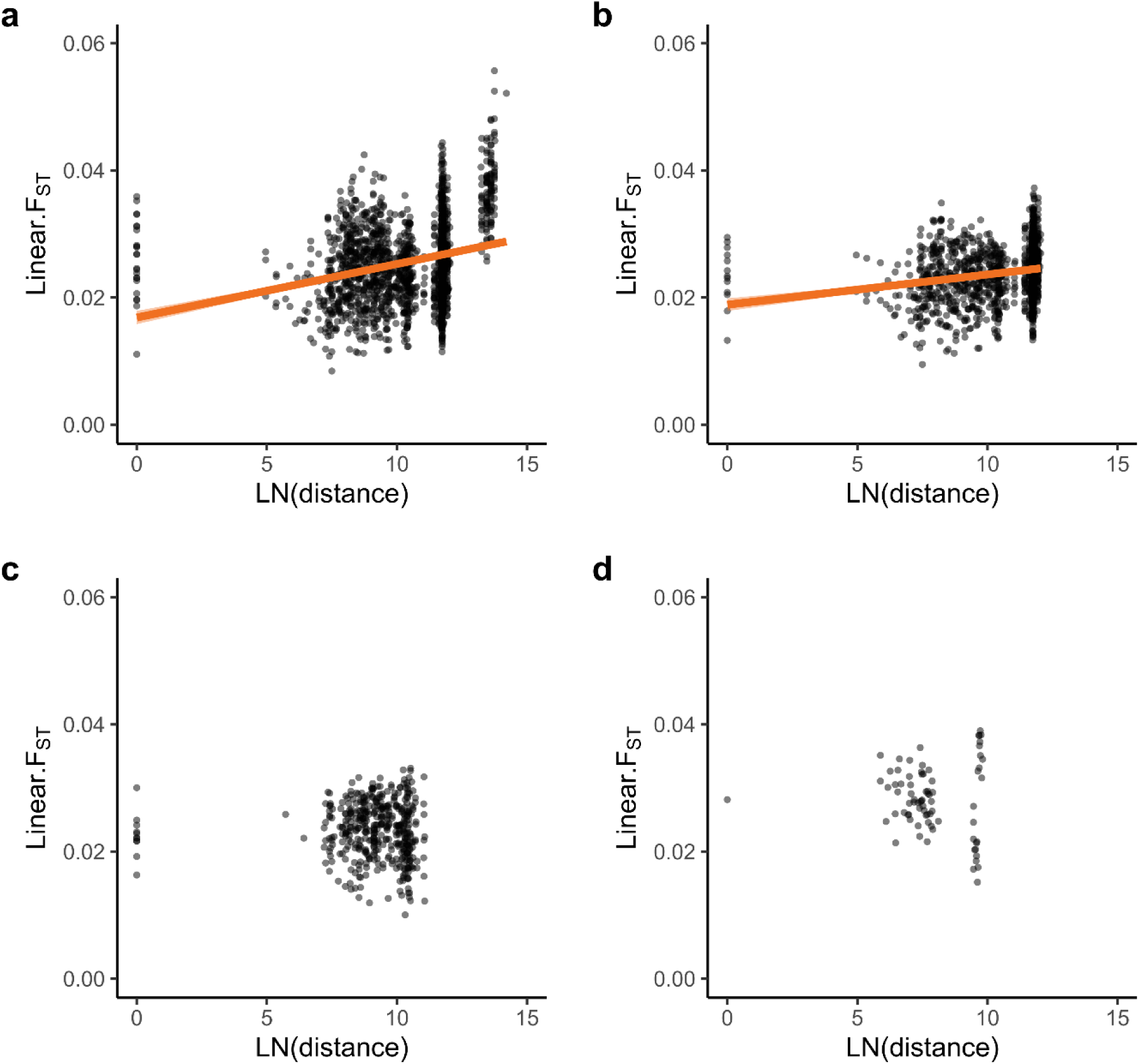
Analysis of isolation by distance. (a) East coast pools. (b) NSW pools. (c) Newcastle pools. (d) Sydney pools with Duck River removed. Including Duck River did not change the plot shape or significance level. Significant Mantel tests are indicted by orange regression lines (in both cases, Mantel R > 0.2, P < 0.0001).

### Variation within individual pools

Heterozygosity and Tajima’s D both varied across Australia (Fig. 5). Overall, the highest heterozygosities were observed in NSW, and the lowest were in WA (Fig. 5a), with the highest values 52% higher than the lowest. Within NSW, Sydney pools had slightly higher heterozygosity (0.00816) than Newcastle (0.00790) (Welch’s t-Test: t = 2.07; d.f. = 42; p = 0.04). Tajima’s D was consistently negative across all regions except WA where scores were closer to zero (Fig. 5b). Heterozygosity and Tajima’s D were generally consistent among sampling sites within each region (Fig. 5 c, d). Geographical interpolation of the east coast scores via exponential kriging clarified these patterns further, highlighting Sydney as the region with highest heterozygosity and most negative Tajima’s D (Fig. 6), though outside of WA variation in Tajima’s D was low.

**Fig. 5.**
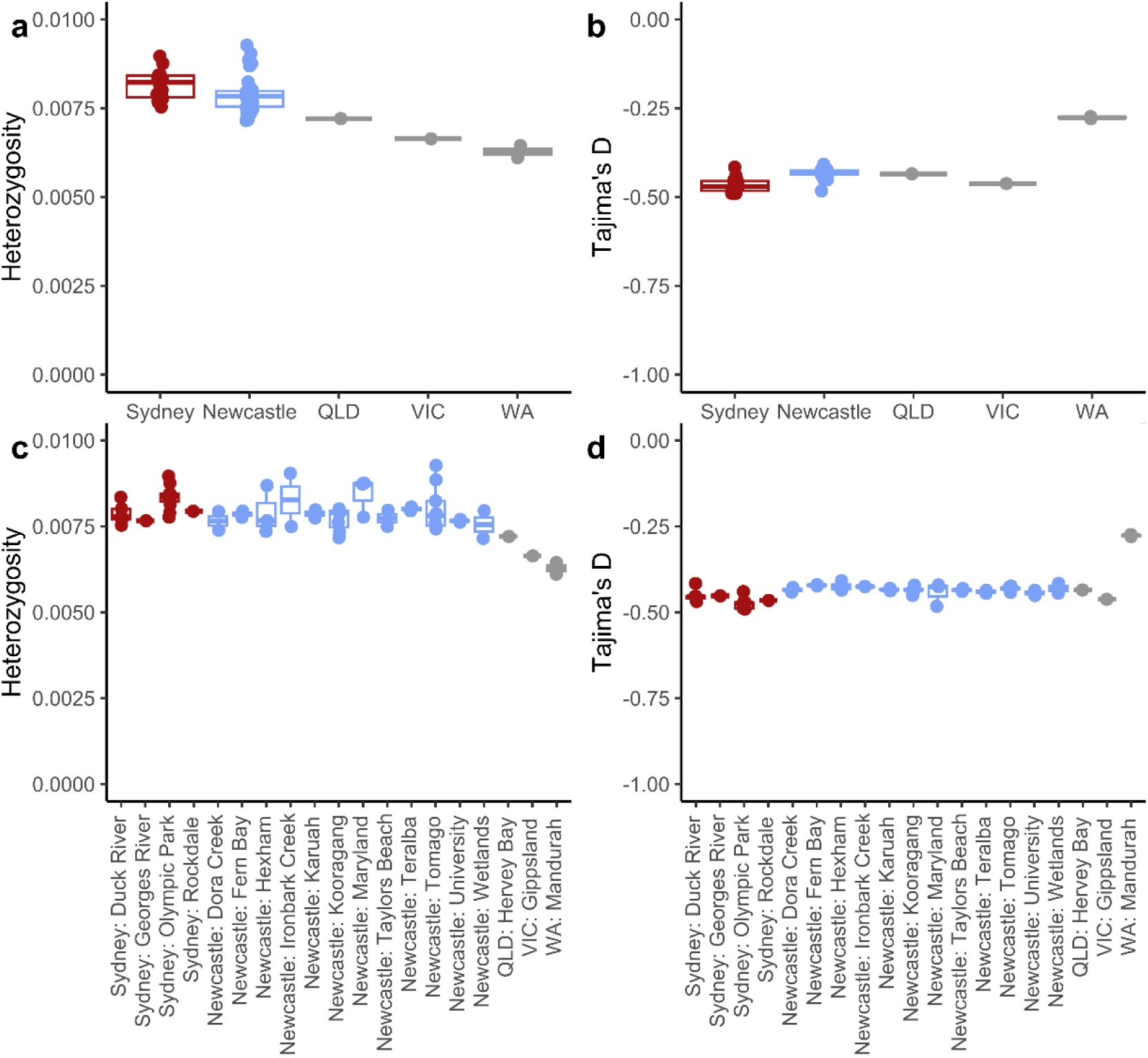
Heterozygosity and Tajima’s D of all pools, aggregated across regions (a, b) or sampling locations (c, d).

**Fig. 6.**
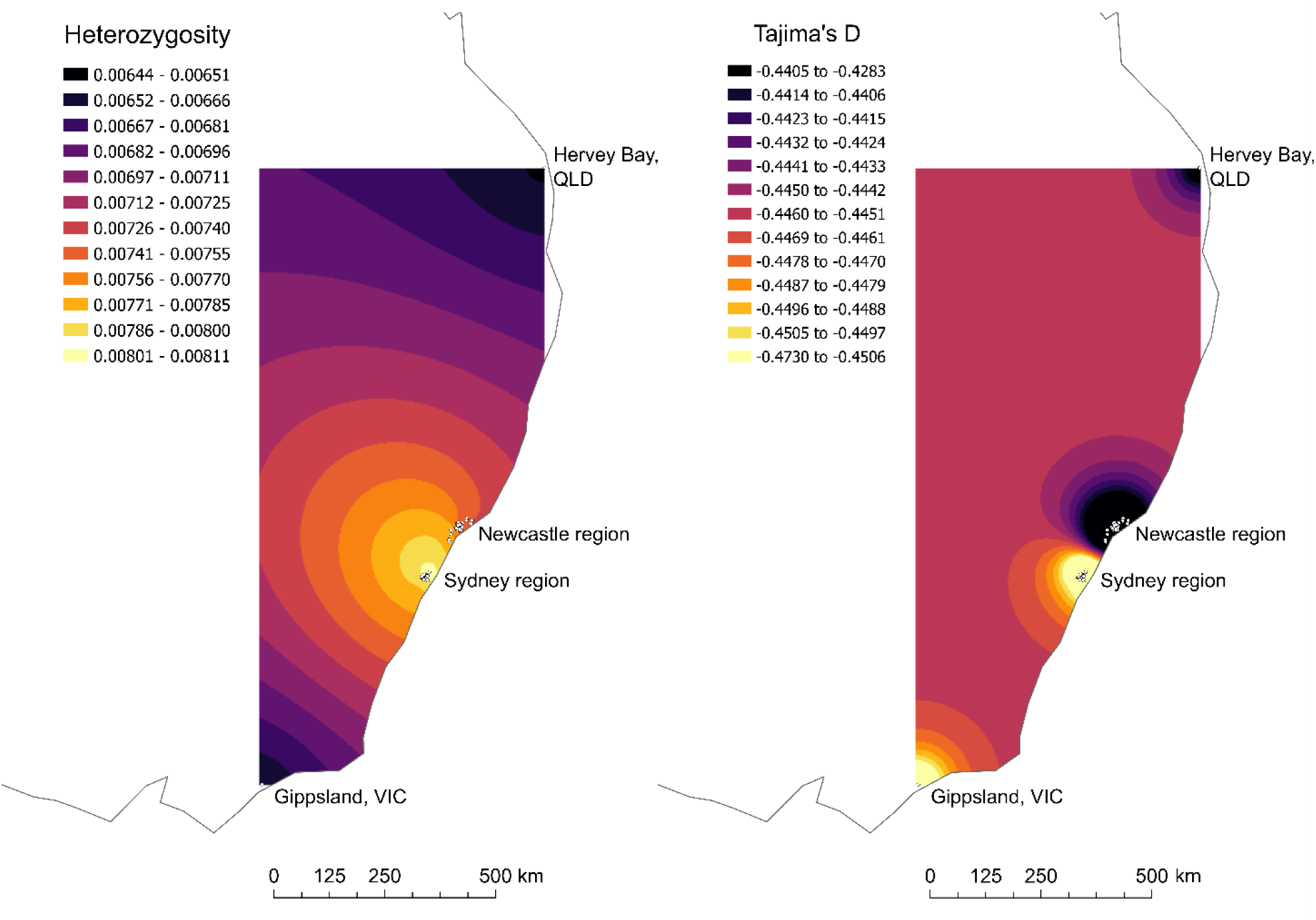
Heterozygosity and Tajima’s D based on pools from the east coast, interpolated across space.

We estimated long-term effective population size in Newcastle and Sydney regions by combining reads from across all 18 Sydney pools and all 38 Newcastle pools into ‘superpools’ for each region. This gave estimates of π of 0.00923 (Newcastle) and 0.00945 (Sydney). Using a recent mutation rate estimate for *Ae. aegypti* (4.85e^-9^; Rose et al. 2023), we estimate long-term effective population sizes of 476,605 (Newcastle) and 487,217 (Sydney). When we consider the broad confidence intervals from an earlier *D. melanogaster* estimate (1e^-9^ – 6.1e^-9^, Keightley et al. 2014), we obtain ranges of 378,940 – 2,311,534 (Newcastle) and 387,377 – 2,363,002 (Sydney).

## Discussion

Our analysis of spatial and temporal genetic structure uncovered several surprising patterns. The first of these was that there was no genetic structure across the Newcastle region. No continuous isolation by distance patterns were observed at ∼60 km scales and the most distant Newcastle pools were no more differentiated than pools from the same location. These results are unlikely to be limited by low analytical power, as the dataset was sufficient to clearly differentiate all the sampling regions as well as temporally stable structuring around Duck River in Sydney. Instead, these results point to a very high level of mobility in *Ae. vigilax*, supporting previous investigations that found large flight ranges in this species (Chapman et al. 1999; Webb and Russell 2019; Johnson et al. 2020). *Aedes vigilax* has been recorded dispersing ∼50 km to offshore islands, possibly by wind (Marks 1969); while rare, long-distance dispersal occurs in many or most mosquitoes, our results may reflect a higher frequency of dispersal at this scale in *Ae. vigilax* than in taxa like *Ae. aegypti* and *Ae. albopictus* where isolation by distance is observed at scales of < 10 km (Schmidt et al. 2017b, 2023a; Yeo et al. 2023). It is worth noting that the preferred habitat of *Ae. vigilax*, tidally influenced saltmarsh and mangrove communities, is widely distributed across the greater Newcastle region. There are extensive areas of estuarine wetland around the Hunter River that produce abundant populations of *Ae. vigilax*. Similarly productive habitats exist to the north of Newcastle, within Port Stephens Bay and the Karuah River, and to the south of Newcastle, around Lake Macquarie. Notwithstanding the potentially high capacity of *Ae. vigilax* to disperse from immature habitats, the widespread abundance of suitable habitat might also have been expected to produce local genetic structure across populations.

Given the apparent panmixia across Newcastle region, it was particularly surprising to observe temporally stable spatial structure between the Duck River and Sydney Olympic Park regions. These were each sampled across two years (2021 and 2022), corresponding to around 6–8 *Ae. vigilax* generations. This structure suggests that recruitment in Duck River was mostly local over this 12-month period, in contrast to recruitment in other locations in Newcastle and Sydney. An alternative hypothesis is that structure in Duck River reflects recent bottlenecks, inbreeding, or strong natural selection over a handful of generations, but this can be rejected by the lack of any difference in heterozygosity or Tajima’s D compared to other Sydney pools. It is unclear why the sample of *Ae. vigilax* from Rockdale was less differentiated from Duck River than the other Sydney samples, despite Rockdale being relatively isolated from the wetlands along the Parramatta River and the Georges River (Sahar et al. 2021).

The drivers of this spatial structure are unknown. Duck River samples were collected only ∼2 km from the Sydney Olympic Park samples. One hypothesis is that dispersal of *Ae. vigilax* from adjacent wetlands into Duck River has been reduced due to mosquito control activities in Sydney Olympic Park over the last ∼20 years (Webb and Russell 1999). The primary mosquito control agent used in these wetlands has been *Bacillus thuringiensis israelensis* (Bti) but there is no evidence that *Ae. vigilax* has become resistant to this control agent to drive mosquito dynamics at fine spatial scales (e.g. Hancock et al. 2022). However, spatially heterogeneous applications of insecticides could still suppress gene flow between areas. A second hypothesis is that Duck River is surrounded by highly urbanised/industrialised land that may operate as an effective dispersal barrier (Fig 3b). Human constructions such as roads are known to limit mosquito movement (Schmidt et al. 2023a), and may be influencing dispersal between Duck River and nearby sites to the east. Local authorities have reported that complaints about mosquitoes from residents and businesses in this residential/industrial region are fewer than expected based on the high mosquito abundance around Duck River (Claflin and Webb 2017), in support of this hypothesis. A final hypothesis is that habitat around Sydney Olympic Park is less suitable for breeding due to low salinity. *Aedes vigilax* is most commonly associated with estuarine habitats, although it has also been recorded from local freshwater environments (Hanford et al. 2020). The presence of *Ae. vigilax* in water with low salinity levels recorded in Sydney Olympc Park (0.18 – 0.68 ppt; Hanford et al. 2020) is unusual. Salinity levels of other sampling points would be mostly higher, including at Duck River (0.06 – 28.1 ppt; (Applied Ecology Pty Limited et al. 2012)). Low salinity could interrupt gene flow across western Sydney.

Despite some local structure and stronger differentiation of the WA population from the east coast populations, our results do not support an initial subdivision of west and east coast populations with subsequent introgression as proposed based on COI data (Puslednik et al. 2012). We appreciate that WA may represent an ancient range expansion rather than recent human transportation, consistent with attenuated variation in WA compared with east coast populations (Puslednik et al. 2012). However, our read alignment rates and levels of genetic structure do not support the notion that *Ae. vigilax* from WA represent a cryptic taxon rather than a highly differentiated population. We also did not find close associations between WA and any east coast populations as might be expected when differentiated clades occur in a region. Our results thus confirm that all populations sampled represent conspecifics, in contrast to cryptic taxa as noted in other mosquitoes (Endersby et al. 2013; Small et al. 2023). Overall patterns of heterozygosity point to an ancestral origin of Australian *Ae. vigilax* in NSW, and populations in QLD and VIC may represent range expansions or native range populations that have been subject to moderate bottlenecks that have not occurred in NSW populations. Effective population sizes of *Ae. vigilax* in NSW are comparable to unsuppressed and non-invasive populations of other *Aedes* taxa such as *Ae. aegypti formosus* (Rose et al. 2023).

### Implications for control of *Aedes vigilax* with *Wolbachia* in Newcastle region

Urban environments like Sydney and Newcastle present unique challenges for the management of arboviral vectors. While most disease transmission tends to takes place in these environments (Duval et al. 2023), it is not well understood how they affect vector biology (Schmidt 2024). The “Anthropogenically induced adaptation to invade” hypothesis proposes that urban environments share characteristics that are homogenous across geography, so that local adaptation to one urban environment serves to adapt the pest to other urban environments (Hufbauer et al. 2012). However, recent research into urban pest evolution has identified considerable fine-scale heterogeneity within urban environments, such as in the distribution of insecticide resistance mutations (Hancock et al. 2022). Urban environmental features may restrict mosquito dispersal (Jasper et al. 2019; Schmidt et al. 2023a) or may facilitate it (Schmidt et al. 2017b; Yeo et al. 2023). These patterns will influence the success of urban pest control programs, including whether strategies will be portable from one region to another, and the appropriate spatial scale over which control programs should operate (Schmidt 2024).

Understanding what our results might mean for any putative *Wolbachia* releases in the Newcastle region requires separate consideration of the two key methods of *Wolbachia*-based control: suppression releases or replacement releases. Suppression releases aim to reduce mosquito population numbers via the release of large numbers of laboratory reared, *Wolbachia*-infected males, which sterilize wild, uninfected females that mate with them. Large-scale releases have been effectively deployed against *Ae. albopictus* in China (Zheng et al. 2019) and against *Ae. aegypti* in Singapore (Bansal et al. 2024), with successful pilot releases in Australia against *Ae. aegypti* as well (Beebe et al. 2021). Following suppression, areas will be recolonised by mosquitoes dispersing into the area, making suppression strategies most effective in isolated populations where recolonisation is slower (Beebe et al. 2021). The absence of genetic structure in *Ae. vigilax* across the Newcastle region suggests that local suppression in subsections of this ∼60 km region will be difficult due to high dispersal rates and require a very large number of mosquitoes to be produced given our estimated effective population sizes of 378,940 – 2,311,534 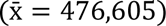 in the region. Operational barriers to distributing infected males effectively will also exist given the extensive and inaccessible wetlands in the region, something which is less problematical in the suppression releases undertaken for other *Aedes* species that have focussed on urbanised areas.

Instead of suppressing a population, replacement releases aim to drive *Wolbachia* into the population through combined releases of male and female mosquitoes. Once local frequencies are sufficiently high, *Wolbachia* will increase in frequency towards fixation as females infected with *Wolbachia* will have a fitness advantage relative to those without it (Hoffmann and Turelli 1997). While replacement releases do not reduce mosquito numbers, many *Wolbachia* strains can strongly reduce virus transmission by infected mosquitoes, which has led to dramatic reductions in dengue transmission in Indonesia (Utarini et al. 2021) and Malaysia (Nazni et al. 2019) and has significantly reduced the risk of local transmission of dengue viruses in Australia (Ryan et al. 2020). If a *Wolbachia* transinfection in *Ae. vigilax* can reduce transmission of arboviruses such as RRV and BFV, it may provide the basis for replacement releases. The large population size and high gene flow in *Ae. vigilax* indicate that initial release numbers may have to be large to establish the infection at a sufficient frequency (e.g. >0.35, Schmidt et al., 2017a); releasing *Wolbachia*-infected mosquito eggs into saltmarshes could represent an effective method to scale such an intervention. Once established, a large population size and high gene flow should serve to maintain the *Wolbachia* infection across a wide area; any local decreases in population numbers such as observed over winter may be countered by *Wolbachia*-infected migrants from an invaded region. Accordingly, we propose that replacement releases may be worth targeting in the Newcastle region if sufficient resources are available to ensure initial establishment of *Wolbachia* across the region.

## Data Availability

All sequence data will be made publicly accessible on the NCBI SRA by time of publication.

## Notes

### Competing Interest Statement

The authors have declared no competing interest.

