## Supplementary material for "Populations of the Australian saltmarsh mosquito *Aedes vigilax* vary between panmixia and temporally stable local genetic structure": Figure S1

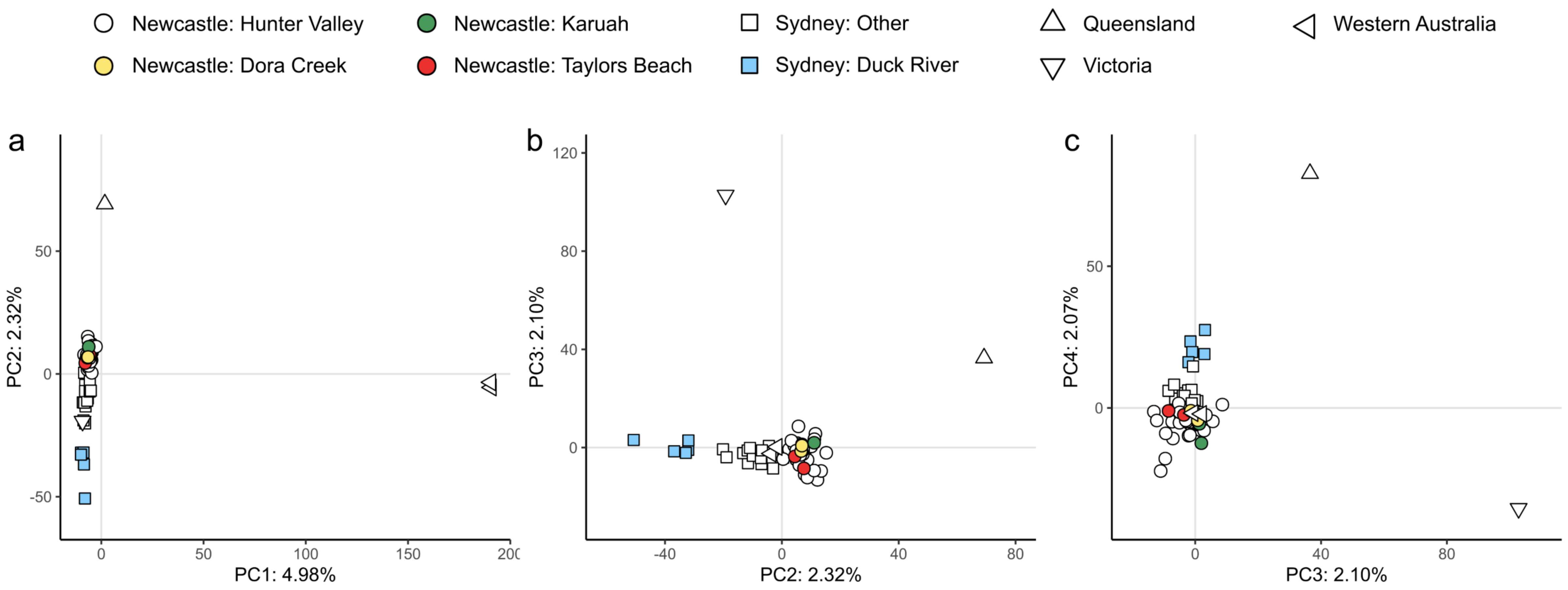


Figure S1. PCA results for all 60 pools. Triangles indicate sampling sites outside of NSW. Coloured symbols indicate peripheral populations around Newcastle and the Duck River population in Sydney. (a) PCs 1 and 2. (b) PCs 2 and 3. (c) PCs 3 and 4.
